# Gain-of-function genetic screens in human cells identify SLC transporters overcoming environmental nutrient restrictions

**DOI:** 10.1101/2022.04.03.486870

**Authors:** Manuele Rebsamen, Enrico Girardi, Vitaly Sedlyarov, Stefania Scorzoni, Konstantinos Papakostas, Manuela Vollert, Justyna Konecka, Bettina Guertl, Kristaps Klavins, Tabea Wiedmer, Giulio Superti-Furga

**Affiliations:** CeMM Research Center for Molecular Medicine of the Austrian Academy of Sciences, 1090 Vienna, Austria; Department of Biochemistry, University of Lausanne, 1066 Epalinges, Switzerland; Center for Physiology and Pharmacology, Medical University of Vienna, 1090 Vienna, Austria

**Keywords:** solute carriers, transporter, amino acids, nutrient limitation

## Abstract

Solute carrier (SLC) transporters control fluxes of nutrients and metabolites across membranes and thereby represent a critical interface between the microenvironment and cellular and subcellular metabolism. Because of substantial functional overlap, the interplay and relative contributions of members of this family in response to environmental stresses remain poorly elucidated. In order to infer functional relationships between SLCs and metabolites, we developed a strategy to identify human SLCs able to sustain cell viability and proliferation under growth-limiting concentrations of essential nutrients. One-by-one depletion of 13 amino acids required for cell proliferation enabled gain-of-function genetic screens using a SLC-focused CRISPR/Cas9-based transcriptional activation approach to uncover transporters relieving cells from the growth-limiting metabolic bottleneck. We identified the cationic amino acid transporter SLC7A3 as a gene that, when upregulated, overcame low availability of arginine and lysine by increasing their uptake. SLC7A5 (LAT1), on the other hand, was able to sustain cellular fitness upon deprivation of several neutral amino acids. A genome-wide screen identified SLC7A3 as the single main gene product able to rescue cell survival in the limiting arginine conditions tested, demonstrating the potentially decisive role of transporters in overcoming nutrient limitations. Moreover, we identified metabolic compensation mediated by the glutamate/aspartate transporters SLC1A2 and SLC1A3 under glutamine-limiting conditions. Overall, this gain-of-function approach using human cells led to the definition of functional transporter-nutrient relationships and revealed that upregulation of transport activity may be sufficient to overcome environmental metabolic restrictions.

## Introduction

A major cellular metabolic requirement is the ability to respond and adapt to the external environment in order to obtain sufficient amounts of the building blocks and sources of energy required for growth and proliferation. When the concentration of a critical nutrient decreases below a certain threshold due to lack of availability or increased consumption, transcriptional and metabolic programs are triggered to limit anabolic and promote catabolic processes via expression of metabolic enzymes or induction of autophagy and macropinocytosis^1–4^. Limiting nutrient availability in the extracellular microenvironment has been shown to deeply affect cell growth, activity and proliferation in both normal and pathological contexts, including development^5^, cell differentiation^6,7^, immune responses^8^ and cancer microenvironment^3,9^. Amino acids in particular represent ∼60% of the dry mass of a cell^2^ and a major source of both biomass and energy for mammalian cells^10^. Importantly, several amino acids, i.e. histidine, isoleucine, leucine, lysine, methionine, phenylalanine, threonine, tryptophan and valine (“essential” amino acids), cannot be directly synthetized by cells and have therefore to be imported from the environment. This requires the action of transmembrane transporters located either at the plasma membrane or, in case of uptake via macropinocytosis or receptor-mediated endocytosis, in the endolysosomal compartment. Similarly, upon starvation, cells induce autophagy to restore nutrient and amino acid levels, and transporters in the autophagolysosome mediate their transfer to cytoplasm^2,11^. Limiting concentrations of amino acids have been reported as metabolic liabilities of cancer cells, including glutamine^12^ and cysteine^13^ depletion in several solid cancers, asparagine in acute lymphoblastic leukemia^14,15^ and arginine in argininosuccinate synthetase-deficient malignancies^15,16^. Accordingly, reduction of the expression of amino acid transporters has been shown to affect cancer cell growth^17–19^ and mouse cancer models showed a tumor-intrinsic dependency on glucose and amino acids transporters^20–23^. Despite these findings, it is not clear if changes in expression of individual plasma membrane transporters alone would be sufficient to relieve cells from metabolic bottlenecks caused by limiting amount of nutrients.

Solute carriers represent the largest family of human transporters, counting more than 450 genes grouped in sub-families based on sequence similarity^24,25^. The fact that a cell expresses at any given time a set of 150-250 solute carriers^26,27^, often with overlapping substrate specificities, results in a high level of functional complexity which hampers efforts to define the individual roles of each protein in nutrient uptake and cell growth. Moreover, cells are known to rearrange the repertoire of expressed transporters upon environmental challenges^28^ and therapeutic treatments^29^.

Forward genetics approaches have been particularly successful in assigning a gene to specific biological processes and pathways, both at the genome-wide level and with more focused approaches^30–33^. Most of the studies published so far focus on loss-of-function (l.o.f.) approaches, as these tend to give binary, easily interpretable, outputs (i.e. wt vs ko phenotypes). However, the availability of CRISPR/Cas9-based approaches has recently expanded the range of readouts and phenotypes that can be explored with large, unbiased gain-of-function (g.o.f.) screens via upregulation of genes from their endogenous loci^34–36^. Importantly, g.o.f. approaches have the advantage of overcoming the issue of redundancy between genes and provide a ranking of the genes that, when overexpressed, confer a selective advantage in a given situation. This aspect is particularly attractive in the context of solute carriers, which have a high degree of substrate and functional redundancy among them^37^.

Here we present the development of a g.o.f. genetic screening approach that allowed us to systematically identify key solute carriers able to overcome nutrient limitation in intact human cells, where the transporters are within natural membranes, with their endogenous modifications and partners. By applying this approach to assess different transporter-based responses to limiting concentrations of 13 amino acids required in our cellular experimental system, we identified SLC7A3 (CAT3), SLC1A2/A3 (EEAT2/1) and SLC7A5 (LAT1) as the transporters able to rescue cells from low availability of arginine/lysine, glutamine and histidine/tyrosine respectively.

## Results

### Amino acid transporters are upregulated upon single amino acid starvation

In order to determine the cellular programs activated by cells upon nutrient starvation and the corresponding regulation of SLCs expression, we generated a transcriptional profile of HEK293T and HeLa cells upon removal of a specific nutrient. HEK293T cells were incubated for 16h in defined media containing dialyzed serum (MWCO 10k Da) and lacking a specific amino acid, vitamins or glucose (Suppl Table 1)^37^. As amino acid starvation impairs mTORC1 activity, we also treated cells with the mTOR inhibitor torin 1^38^. To evaluate general and cell type-specific effects, responses to single amino acid depletion were also assessed in HeLa cells. Transcript abundance was determined by 3’ mRNA sequencing and enrichment was calculated by comparing the samples under nutrient limiting conditions to the fully reconstituted media. We observed large transcriptional changes in most samples, with patterns consistent across the two cell lines (Fig 1a-b, Suppl Fig 1a-b). One exception was observed upon deprivation of serine or glycine, which showed only marginal changes, likely as a result of the cell’s ability to readily interconvert these two amino acids^39,40^. Interestingly, absence of vitamins, as tested in HEK293T cells, did not result in large transcriptional changes in this experimental setup. Moreover, while we observed comparable changes in most of the other single amino acid depleted conditions, methionine depletion appeared to induce a distinct transcriptional response, similar to previous reports^41^. To identify the transcriptional programs induced by nutrient limitation, we calculated the enrichment of transcription factor target genes in the tested samples based on the TRRUST dataset^42^. We observed the ATF4 transcription factor program as the top enriched among most amino acid-depleted samples (Fig 1c, Suppl Fig 1c), consistent with the role of this transcription factor downstream of the amino acid sensor GCN2 in activating the amino acid response (AAR)^28^. Accordingly, we detected extensive changes in the mRNA abundance of amino acid transporters under conditions of limiting amino acid compared to fully reconstituted media (Fig 1d, Suppl Fig 1d). More than 60 amino acid transporters have been described, comprising members of the SLC1, SLC7, SLC36, SLC38 and SLC43 subfamilies^43^. Interestingly, a subset of transporters appeared to be highly upregulated in most aa-limiting conditions, regardless of the amino acid removed from the media and consistent with the induction of AAR (Fig 1d, Suppl Fig 1d). Within this set of SLCs, we observed several amino acid transporters, including members of the SLC1 (SLC1A4 and SLC1A5), SLC7 (SLC7A1, SLC7A3, SLC7A5, SLC7A11 and the associated SLC3A2 heavy chain) and SLC38 families (SLC38A1, SLC38A2). Of note, several of these transporters are controlled by ATF4, including SLC1A5 (ASCT2), SLC7A1 (CAT1) and SLC7A11 (xCT), and have been reported to play a role in maintaining amino acid homeostasis^18,44^. These results confirmed that a profound adaptation of transporter expression is a prominent phenomenon in the responses induced to overcome nutrient starvation, and amino acid limitation in particular. Most importantly, this raises the question whether the observed changes in transporter expression are sufficient to overcome nutrient limitations and which of these SLCs may contribute most to restore cellular fitness in each of these conditions.

**Figure 1:**
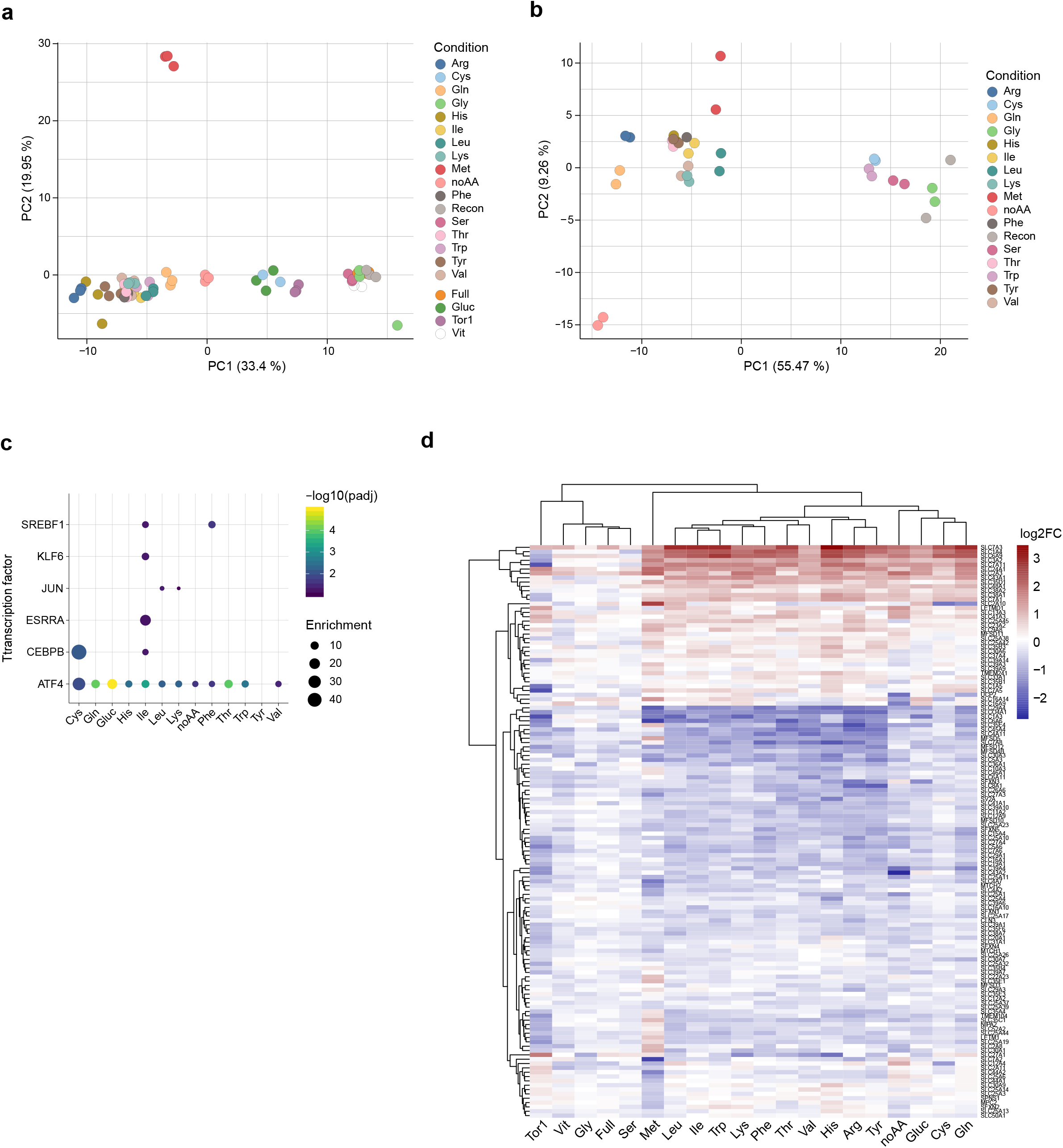
**a**. PCA plot of the transcriptomics profiles of HEK293T cells upon specific nutrient deprivation conditions or Torin-1 treatment for 16 hours. Replicates for each condition are shown with the same color code. **b**. PCA plot of the transcriptomics profiles of HeLa cells upon specific nutrient deprivation conditions for 16 hours. Replicates for each condition are shown with the same color code. **c**. Enrichment analysis for target genes of transcription factors in the HEK293T samples, as determined by TRRUST v2. **d**. Heatmap showing differential expression of SLC transporters in the listed conditions compared to reconstituted media in HEK293T cells.

### Generation of a SLC-focused gain-of-function library and development of nutrient-dependent cellular fitness screens

In order to systematically interrogate the SLC superfamily and identify specific transporters involved in the response to nutrient limitation, we generated a gain-of-function lentiviral-based CRISPR/Cas9 library targeting 388 human SLCs (SLC-SAM library, Fig 2a, Suppl Fig 2a, Suppl Table 2). The library was based on the Synergistic Activation Mediator (SAM) approach^35^, which has been shown to be one of the most effective CRISPR-based approaches developed for transcriptional activation^34^. This approach relies on a nuclease-inactive Cas9 fused to a transcriptional activator domain (VP64) and MS2-based recruitment of p65-HSF1 to induce sgRNA-driven overexpression of a given gene from its endogenous promoter. Up to six sgRNAs per SLC transcript were selected. Representation within the library was determined by next-generation-sequencing (NGS, Suppl Fig 2b).

**Figure 2:**
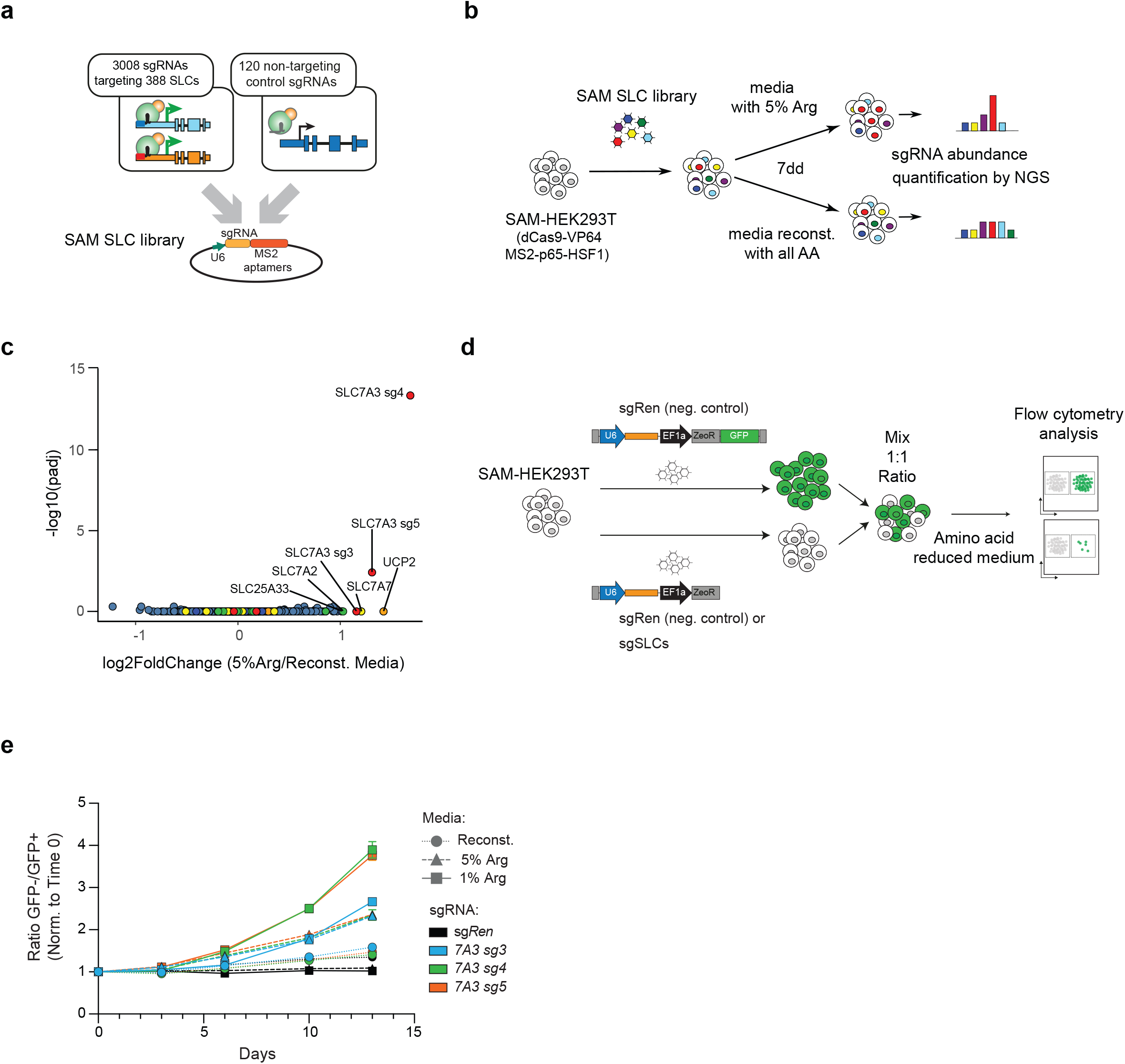
**a**. Schematic representations of the SLC SAM library used in this study. **b**. Experimental setup for the screening for SLCs able to overcome Arginine limitations in the extracellular media **c**. Volcano plot showing enrichment of sgRNAs in arginine-limiting conditions against fully reconstituted media after 7 days of treatment. Log2 fold changes and adjusted p-values were calculated using DESeq and GSEA from two replicate experiments. **d**. Schematic representation of the color competition assay (CCA) approach. **e**. Time-dependent enrichment of SAM-HEK239T cells carrying sgRNAs targeting SLC7A3 in limiting arginine media concentrations. Data shown derived from one experiment performed in three replicates. Results are representative of two independent experiments.

In order to perform genetic screens, we set out to identify conditions in which reduced levels of specific amino acids would confer a sufficient selective pressure. Thus, HEK293T were grown in amino acid-free media reconstituted with all amino acids at their original concentration with the exception of one particular amino acid. We chose to test the set-up using arginine, which was supplemented at 0.02 mM, a concentration corresponding to 5% of the one present in normal, rich media (0.4 mM). Media was further supplemented with dialyzed serum. As expected, this resulted in reduced proliferation and cell death (Suppl Fig 2c), thereby conferring the selection pressure required for the planned g.o.f screen. We therefore selected this limiting arginine condition to test whether the SLC-SAM library we generated allowed for the unbiased identification of transporters able to restore cell proliferation under nutrient starvation. We infected HEK293T stably expressing the SAM transcriptional activation machinery (SAM-HEK293T) with the SLC-SAM library and, after antibiotic selection, switched to media containing either 5% arginine or fully reconstituted. After 7 days in culture, the remaining cells were collected and the composition of the library determined by sequencing (Fig 2b, Suppl Table 3). Enrichment analysis of the samples grown in low-arginine versus the fully reconstituted conditions showed a strong and statistically significant enrichment for sgRNAs targeting the sodium-independent cationic amino acid transporter SLC7A3 (CAT3) (Fig 2c). This protein is able to transport arginine, as well as lysine and ornithine, with K*m* in the micromolar range^45^, therefore strongly supporting the ability of our approach to identify direct transporter/substrate relationships. Of note, several members of the SLC7 (CAT) subfamily of cationic amino acid transporters were indeed enriched in our transcriptomics analysis (Fig 1d).

### SLC7A3 upregulation overcomes arginine limitation

To validate the previous results, we next generated cells individually expressing the three enriched sgRNAs targeting the *SLC7A3* (sg*SLC7A3*) promoter or a control sgRNA encoding a *Renilla* sequence (sg*Ren*). All *SLC7A3* sgRNAs led to an upregulation of *SLC7A3* transcription as confirmed by q-PCR (Suppl Fig 2d). To functionally assess whether increased levels of SLC7A3 confer resistance to reduced levels of arginine, we used color competition assay (CCA) (Fig 2d). sg*SLC7A3*- or control sg*Ren*-expressing cells were seeded in a 1:1 ratio with cells co-expressing control sg*Ren* and GFP, allowing us to monitor differential growth over time by flow cytometry. Validating the results obtained in the gain-of-function screen, *SLC7A3*-overexpressing cells showed increased fitness compared to control cells when cultured in low arginine media, while no difference was observed in fully reconstituted media or when sg*Ren*-expressing cells were used (Fig 2e). Interestingly, the competitive advantage of sg*SLC7A3*-bearing cells was stronger at the lowest concentration of arginine and enrichment increased over culture time and cell replating. Together, these data support the validity of our gain-of-function approach to identify SLCs sustaining cell fitness upon nutrient deprivation and highlight SLC7A3 upregulation as a strategy to resist arginine scarcity.

### A systematic screen for transporters overcoming extracellular amino acid limitation

Encouraged by these results, we extended this approach to systematically screen for SLCs sustaining cellular fitness under limiting concentrations of other amino acids. We therefore tested the effect on cell fitness of selectively reducing the concentration of each of the 15 amino acids present in DMEM culture media and observed a strong impairment of cell viability for all amino acids with the exception of glycine or serine, in line with the lack of significant transcriptional responses observed upon depletion of these two amino acids (Fig 1 and Suppl Fig 1). Based on the arginine and SLC7A3 validation results (Fig 2e), we selected conditions with reduced concentration of amino acids imposing a strong selective pressure (between 0.1-2% of the normal DMEM concentration, Suppl Fig 3a) and performed the screen over 14 days including three passages where cells were detached and re-seeded in fresh media (Fig 3a). SAM-HEK293T cells were transduced with the SLC-SAM library and cultivated in media with reduced levels of single amino acids or lacking all amino acids, while cells grown for the same period in media reconstituted with all amino acids served as control (Suppl Tables 4-7). Figure 3b provides an overview of the screening outcome highlighting significantly enriched SLCs for each condition. Confirming the robustness of our approach, we could recapitulate in these settings the specific enrichment of *SLC7A3* sgRNAs in arginine-reduced conditions (Fig 3b-c). Moreover, specific enrichment of *SLC7A3* was also observed in lysine-reduced conditions (Fig. 3b), a second amino acid described to be a substrate for this transporter^45^. Indeed, monitoring *SLC7A3*-sgRNA abundance across all the tested conditions, we observed that three sgRNAs accumulated specifically in Arg/Lys-reduced media while two additional sgRNAs did not, possibly because the latter were less efficient in triggering transcriptional activation (Fig 3d). In addition to SLC7A3, another member of the SLC7 family of amino acid transporters, SLC7A5 (LAT1), was identified in multiple conditions (Fig 3b, Suppl Fig 3b). SLC7A5 belongs to the second subgroup (after the CAT proteins) of the SLC7 family, the HATs (heterodimeric amino acid transporters), which work as obligate heterodimers with SLC3 family members and display differences in term of substrate specificity and mode of transport^45^. These heteromeric amino acid transporters work mostly as exchangers, and SLC7A5/SLC3A2 substrates comprise large neutral amino acids which are translocated in a sodium- and pH-independent manner. SLC7A5 was the most enriched gene in the screens for histidine, tyrosine, phenylalanine and methionine (Fig 3b, Suppl Fig 3b) and, with a less stringent sgRNA cutoff, also with isoleucine and leucine (Suppl Fig 3c), therefore recapitulating most of the known transporter substrates.

**Figure 3:**
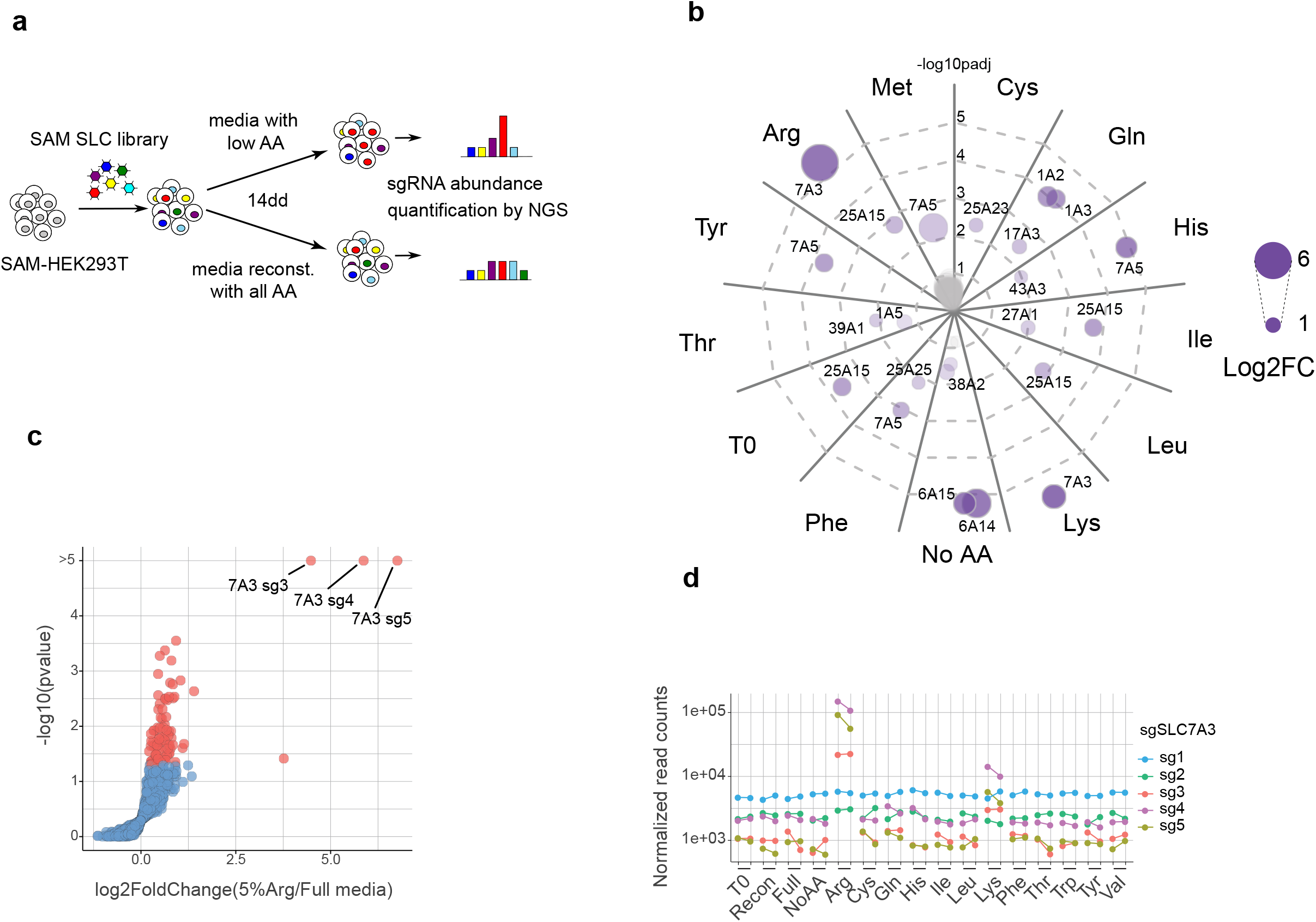
**a**. Experimental setup of the screens for SLCs able to overcome amino acid limitations in the extracellular media. **b**. Overview of the significantly enriched SLCs for each of the tested conditions at FDR< 0.01. The circle size reflects the log2 fold change above the fully reconstituted media, while the distance from the center reflects the statistical significance of the effect observed. **c**. Volcano plot showing the enrichment and statistical significance of each sgRNA upon Arginine starvation. **d**. Plot showing the normalized counts of SLC7A3-targeting sgRNAs across the conditions tested.

Analysing the other screening conditions, we observed enrichment of the aspartate and glutamate transporters of the SLC1 family, SLC1A2 (EAAT2/GLT-1) and SLC1A3 (EAAT1/GLAST-1) upon glutamine starvation (Fig 3b, Suppl Fig 3b-c). This is consistent with recent reports showing that cytosolic aspartate concentrations determine cell survival upon glutamine starvation and that expression of SLC1A3 overcomes the effects of limiting glutamine availability^46,47^. We also detected enrichment for the mitochondrial proton-coupled ornithine and citrulline transporter SLC25A15^48^ in several conditions (including full media and initial, time 0 condition) as a result of sgRNA depletion in the fully reconstituted media condition (Fig 3c). Finally, we observed an enrichment for the two broad-specificity, concentrative amino acid transporters SLC6A14 and SLC6A15^43^ when all amino acids were removed from the culture media, providing therefore conditions where the only amino acid sources were dialyzed serum or possibly dying cells (Fig 3c, Suppl Fig 3b-c). Overall, our g.o.f screen identified key concentrative amino acid transporters able to overcome specific and general amino acid limitation in the extracellular medium.

### Identified SLCs sustain cellular fitness upon starvation of their amino-acid substrates or via metabolic compensation

To validate further the specificity and characterize the role of the identified transporters upon amino acid limitation, we tested several of these SLCs using the above-described color competition assay. sg*SLC7A3*-expressing cells showed growth advantage in media containing low levels of the cationic amino acid arginine and lysine, but not in case of the non-substrate histidine nor in reconstituted media (Fig 4a). We then investigated the role of the other SLC7 member showing enrichment in multiple conditions and confirmed upregulation at the protein level of SLC7A5 by the three top-scoring sgRNAs identified in the screen (Fig 4b-c). Interestingly, sg*SLC7A5*-expressing cells showed a concomitant increase in SLC3A2 protein abundance, which correlated with SLC7A5 induction (Fig 4c). This suggests that upregulated SLC7A5 assembles with its SLC3A2 partner in functional heterodimers and that proteostatic co-regulatory mechanisms control SLC3A2 in response to SLC7A5 levels. In CCA, SLC7A5 transcriptional activation conferred advantage in histidine- and tyrosine-, but not arginine-low media (Fig 4b). These results demonstrated that different SLC7 family members sustain cellular fitness in conditions where cells are starved from one of their amino acid substrates. Finally, transduction of SAM-HEK293T cells with the top three sgRNAs targeting SLC1A2 or SLC1A3 resulted in increased protein levels of the two transporters (Suppl Fig 4a-b) and conferred a modest but consistent selective advantage in low glutamine media (Fig 4d-e), therefore confirming the screen outcome. To further investigate this effect and compare the results obtained by the SAM-mediated upregulation of SLC genes from the endogenous promoter with the ectopic, cDNA-driven overexpression of these same transporters, we established an analogous CCA using HEK293T cells lines expressing codon-optimized versions of SLC1A2 and SLC1A3 as well as GFP under the control of a doxycycline-inducible promoter. Treatment with doxycycline for 24h resulted in a substantial increase of transporter expression (Suppl Fig 4c-e) and immunofluorescence experiments confirmed the plasma membrane localization of the overexpressed SLC1A2/3 proteins (Suppl Fig 4f-g). We then monitored differential growth in CCA by co-culturing parental (empty), SLC1A2, or SLC1A3 overexpressing lines together with GFP expressing cells. Supporting the results obtained with the SAM approach, cells expressing SLC1A2 or SLC1A3 displayed increased fitness specifically in low glutamine, but not in fully reconstituted media, irrespective of the presence of an N-terminal Strep-HA tag (Fig 4f-g). Interestingly, a similar fitness advantage was observed also in media completely devoid of glutamine, supporting the notion that SLC1A2 and SLC1A3 upregulation leads to metabolic compensatory effects and not to direct glutamine uptake. Overall, these data validate the ability of a set of concentrative transporters to overcome amino acid limitations.

**Figure 4:**
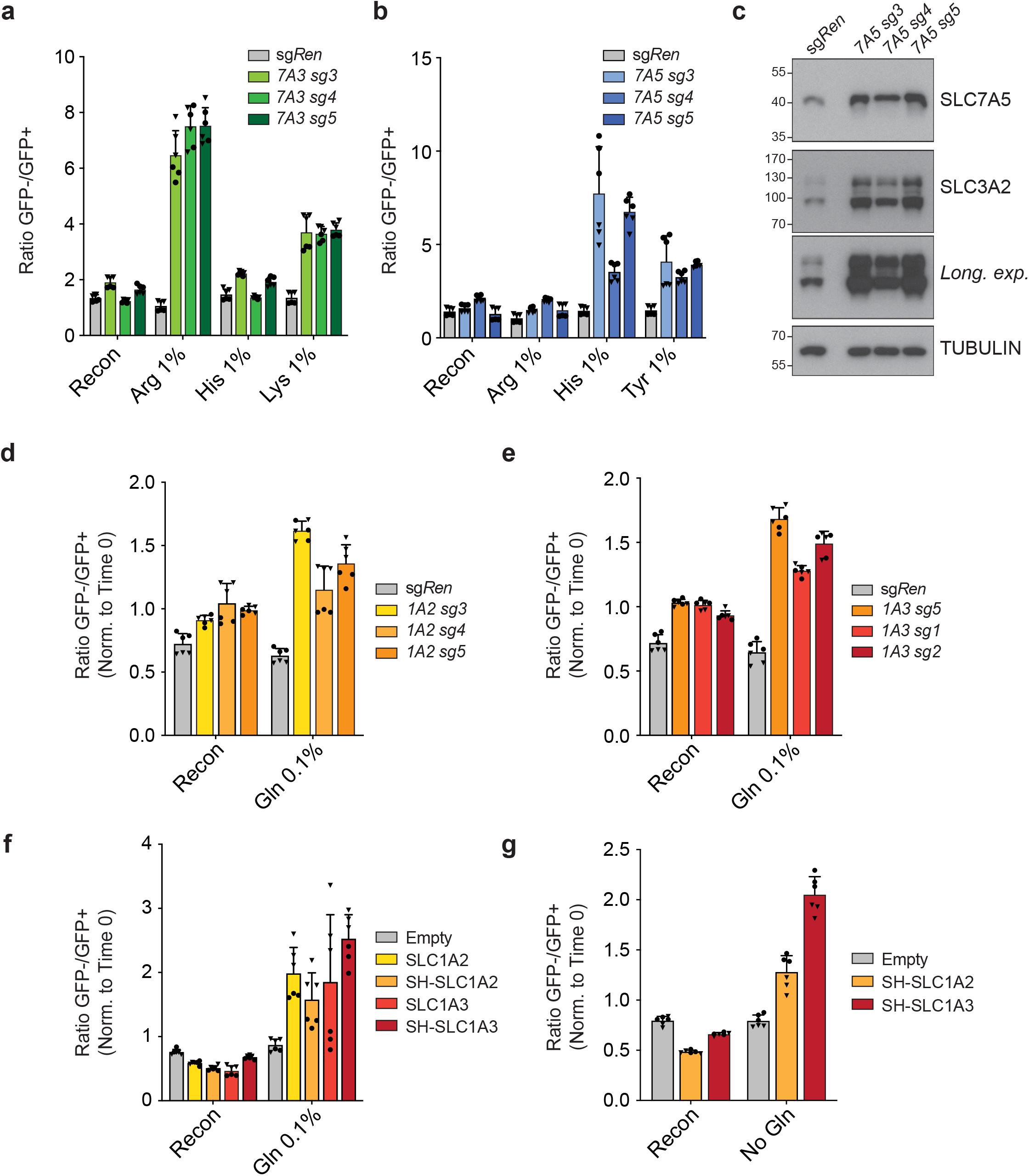
**a**. Color competition assay (CCA) showing enrichment of eGFP-negative SAM-HEK239T cells carrying sgRNA targeting SLC7A3 or sgRen compared to eGFP positive, sgRen carrying control cells in media conditions with limiting concentrations of arginine, lysine but not histidine. Data shown for two independent experiments each performed in triplicates. **b**. CCA showing enrichment of SAM-HEK239T cells carrying sgRNA targeting SLC7A5 in media conditions with limiting concentrations of histidine, tyrosine but not arginine. Data shown for two independent experiments each performed in triplicates. **c**. Western blot showing increased protein levels of SLC7A5 and the accessory chain SLC3A2 in SAM-HEK239T cells carrying sgRNA targeting SLC7A5. **d-e**. CCA showing enrichment of SAM-HEK239T cells carrying sgRNA targeting SLC1A2 (**d**) or SLC1A3 (**e**) in media conditions with limiting concentrations of glutamine. Data shown for two independent experiments each performed in triplicates. **f-g**. CCA showing enrichment of HEK239 Flip-IN cells ectopically expressing untagged or Strep-HA (SH) tagged SLC1A2 or SLC1A3 compared to control HEK239 Flip-IN cells expressing eGFP in media conditions with limiting concentrations (0.1%) (**f**) or absence (**g**) of glutamine. Cells were treated with doxycycline for the duration of the assay. Data shown for two independent experiments each performed in triplicates.

### A genome-wide g.o.f. screen supports SLC7A3 as the prominent gene able to overcome extracellular arginine depletion

A question raised by the SLC-focused approach we developed and validated is how SLC upregulation compares to other possible cellular mechanisms in sustaining cell viability under starvation conditions. The biased nature of the approach focused on SLC transporters, with all its advantage to functionally ascribe transporter-nutrient relationships, does not allow to gauge their relative importance. We thus switched to a genome-wide perspective and performed a g.o.f. screen in arginine-limiting conditions using a library covering all protein-encoding gene products (Fig 5a). We infected SAM-HEK293T cells with a genome wide SAM library carrying 70,290 sgRNAs targeting 23,430 genes^35^ and cultivated them with 1% arginine or fully reconstituted media before collecting samples 14 days later. Comparison of the sgRNA abundances between the arginine-limited and reconstituted media showed a clear enrichment for multiple sgRNAs targeting SLC7A3 (Fig 5b, Suppl Tables 8-9). Accordingly, when analysed at the gene level, SLC7A3 was the single most enriched gene identified, confirming the accuracy of our SLC-focused screens. Importantly, this result suggests that indeed, increased uptake of arginine by this transporter was the most effective mechanism to maintain growth-limiting amino acid homeostasis, at least under these experimental settings (Suppl Table 9). In order to gain a better mechanistic understanding of how SLC7A3 upregulation promotes cell survival and support our hypothesis that this occurs by increasing the uptake of its substrate amino acids, we performed a targeted metabolomic analysis of SLC7A3-overexpressing cells (Fig 5c, Suppl Fig 5a-c, Suppl Table 10). Rewardingly, this revealed that doxycycline-mediated upregulation of SLC7A3 resulted in a striking and specific increase in the intracellular concentration of its substrates arginine, lysine and ornithine^45^. While most of the other metabolites were not affected, we observed an increase also of biogenic amines (symmetric dimethylarginine and alpha-aminoadipic-acid), likely resulting from the elevated cellular levels of arginine and lysine, respectively.

**Figure 5:**
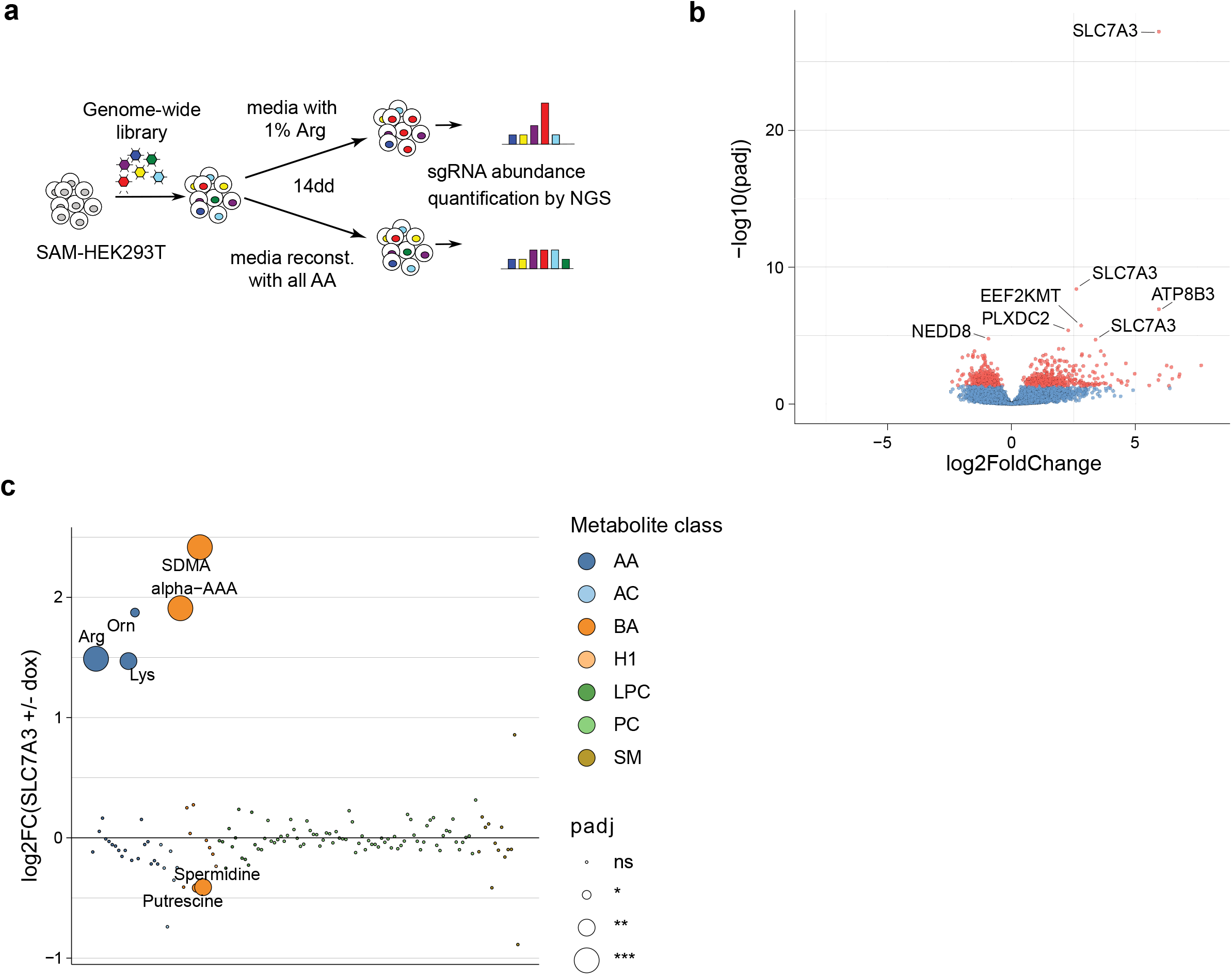
**a**. Schematic representation of the experimental setup for the genome-wide SAM screen upon limiting arginine conditions. **b**. Volcano plot showing sgRNA enrichment between low arginine and fully reconstituted conditions. **c**. Targeted metabolomics analysis of doxycycline-induced HEK293 Jump In cells overexpressing SLC7A3, compared to non-induced cells, shows enrichment for transporter substrates and related compounds. AA. Amino acids; AC, acylcarnitines; BA, biogenic amines; H1, hexoses; LPC, lysophosphoatidylcholine; PC, phosphoatidylcholine; SM, sphingomyelin.

These results highlight the prominent effect of transporter upregulation in supporting the ability of cells to overcome limiting nutrient conditions by increasing their uptake.

## Discussion

The role of cellular metabolic responses to environmental changes and their effect on survival and proliferation in normal and pathological states is increasingly recognized^3,49^. Indeed, metabolic plasticity of human cells is an area of intense research as it is associated with a great variety of processes, ranging from T-cell activation^50^ to metabolic symbiosis within the tumor microenvironment^51^. The mechanisms by which human cells sense nutrient levels and adapt accordingly their metabolism are beginning to be elucidated and typically involve master regulators such as mTOR and AMPK, changes in transcription factors activity, changes in the expression of rate-limiting enzymes and of membrane transporters^2,3^. Transporter (up)regulation is part of the cellular response to nutrient limitation, such as in low glucose^52,53^ and low glutamine^46,47^ conditions. Accordingly, several transporters are part of concerted transcriptional programs initiated by nutrient sensing mechanisms^4,54^. What we wanted to clarify was if transporters are mere executing “workhorses” of metabolic adaptation or also capable, by their own action, to sustain metabolic corrections leading to cell state and cell fate decisions (e.g. growth, no growth). Another recurrent challenge resides in ascribing individual functions to transporters in light of the high level of functional redundancy and plasticity in occurring expression patterns.

The transcriptional responses induced by depletion of individual amino acids in HEK293T and HeLa cells were largely independent of the specific amino acid depleted and included overexpression of a conserved set of amino acid transporters. Several of the well-characterized amino acid transporters families, including the SLC1, SLC7 and SLC38 solute carrier subfamilies, were well represented.

Using a highly scalable SLC-focused CRISPR/Cas9-based approach we developed, we identified both cases of direct uptake of the limiting nutrient by the enriched transporter as well as cases where the transporter overexpression is likely to rescue the nutrient limitation indirectly by providing metabolically related compounds. As examples of the first case, we identified two members of the SLC7 family, SLC7A3 (CAT3) and SLC7A5 (LAT1), as the main transporters able to restore cell fitness upon low extracellular concentration of arginine and lysine, for SLC7A3, and multiple amino acids including histidine, tyrosine and methionine for SLC7A5. We were therefore able to identify, starting from a completely unbiased approach, the known substrates for both transporters, highlighting the potential of this approach for substrate/transporter associations and functional annotation of uncharacterized transporters. Importantly, overexpression of SLC7A3 (CAT3) was confirmed in a genome-wide g.o.f. screen as the top case of single gene overexpression able to overcome arginine limitation. This was quite surprising and revealed a perhaps underestimated role of transporters upon environmental limitations. Exemplifying the second scenario, we identified the aspartate/glutamate transporters SLC1A2 and SLC1A3 as the two solute carriers able to sustain cellular fitness in media containing low or no glutamine. Glutamine is the most abundant free amino acid in cells, exerting a critical energetic role by providing intermediates for the TCA cycle. Recent studies demonstrated that aspartate uptake is able to rescue cells from limiting glutamine conditions^46,47^. Since DMEM media does not contain aspartate and glutamate, it is likely that these may derive from dying cells. Further studies will be needed to investigate this hypothesis.

While these cases refer to specific amino acid conditions, the finding that the broad-substrate amino acid transporters SLC6A14 and SLC6A15 were enriched specifically upon low extracellular concentrations of all amino acids is intriguing. One possibility is that these transporters are particularly energetically expensive as they couple amino acid uptake to the transport of both sodium and chloride ions. Considering the growing interest in SLC6A14^20,21,55^ and amino acid transporters in general as a potential cancer target, these results warrant further studies to understand the effect of environmental conditions on transporters expression.

Solute carriers represent the largest family of transporters in humans and a relatively understudied family, with ∼30% of its members still lacking endogenous substrates despite an increasing recognition of their potential as drug targets and major roles in metabolism^25,37^. The approach described here allows for the systematic interrogation of the solute carrier family to identify the transporter(s) able to rescue nutrient limiting conditions, providing insights into the range and potency of the responses available to cells in challenging environments. Large-scale SLC-focused gain-of-function screens may thus represent a powerful and efficient approach to identify the relevant transporters by their function, and likely contribute to the ongoing efforts of de-orphanization^56^ by overcoming the issue of functional redundancy. An obvious limitation of the strategy concerns those metabolites that are not rate-limiting for growth and would require the development of specific screening read-outs. Another potential shortcoming is the possibility that, when overexpressed, transporters may lose activity or selectivity due to the absence of a partner or a modification.

Despite these limitations, the results presented here provide compelling data to support the hypothesis that upregulation of individual SLCs membrane transporters alone can be sufficient to overcome metabolic bottlenecks. Transporters, alone and in combinations, may thus exert a more central role than generally estimated in governing the metabolic plasticity required for cellular and tissue homeostasis. Because of their role in mediating the cellular needs with the environment, they may be ideally suited to act as conveyors of metabolic integration across tissues. Their dynamic mRNA and protein expression profiles may in future be used to interpret metabolic states of cells and tissues. Altogether, the role in overcoming metabolic bottlenecks increases their attractivity as potential drug targets. In this sense, the approach presented here adds another assay for the identification of targeting chemotherapeutics.

## Author contributions

M.R. and G.S-F conceived the study. E.G. designed, and E.G. and J.K. generated the SLC SAM library. M.R., S.S., K.P. and M.V performed experiments. T.W., B.G and K.K performed metabolomic experiments. V.S. analysed transcriptomics, NGS, and metabolomics data. M.R., E.G and G.S.-F. wrote the manuscript with contributions from all authors.

## Acknowledgements

We would like to acknowledge the members of the Superti-Furga group for critical discussions and suggestions. We thank Johannes W. Bigenzahn for kindly providing critical reagents, Svenja Onstein for experimental support and Ulrich Goldmann for computational support. Part of this work was carried out within the RESOLUTE project. This project has received funding from the Innovative Medicines Initiative 2 Joint Undertaking (JU) under grant agreement No 777372. The JU receives support from the European Union’s Horizon 2020 research and innovation programme and EFPIA. This work was supported by the Austrian Academy of Sciences and the European Research Council (ERC) Advanced Grant 695214 GameofGates.

## Data availability

Genomics datasets are provided in Suppl Tables 3-8. The SLC SAM library is available on Addgene (Addgene 132561). The transcriptomics datasets are deposited at GEO (GSE153034 and GSE199252). Metabolomics data are deposited at MetaboLights (MTBLS4268).

## Methods

### Cell lines and reagents

HEK293T were obtained from ATCC, HeLa were provided by M. Hentze. HEK293 Jump-In and Flp-In TREx cells that allow doxycycline-dependent transgene expression were from Invitrogen. Cell lines were checked for mycoplasma infection by PCR or ELISA. Cell lines were authenticated by STR profiling. Cells were kept in DMEM (Gibco or Sigma) supplemented with 10% (v/v) FBS (Gibco) and antibiotics (100 U/ml penicillin and 100 mg/ml streptomycin, Gibco or Sigma) and shifted to specific nutrient restricted media for the different assay (described below).

### Antibodies

Antibodies used were SLC7A5 (Cell Signaling, #5347), SLC3A2 (Santa Cruz, 376815, E-5), tubulin (Abcam, Ab-7291), HA (Covance, MMS-101P), Actin (Cytoskeleton, AAN01-A), SLC1A2 (Santa Cruz, 365634), SLC1A3 (Cell Signaling, #5684T), HRP-conjugated secondary antibodies (Jackson Immunoresearch).

### Plasmids and cell line generation

SgRNA were cloned into lenti sgRNA(MS2)_zeo backbone (Addgene: 61427). Cloned oligonucleotides, named accordingly to the SLC SAM library ID, were as follows: SLC7A3 (sg3 F caccgGGCTTTGCAAAAGGATTGCG, R aaacCGCAATCCTTTTGCAAAGCCc ; sg4 F caccgTGAGGATGGGACGCAGTCTC, R aaacGAGACTGCGTCCCATCCTCAc; sg5 F caccgTAGCGAGGAGGATTGGGGGT, R aaacACCCCCAATCCTCCTCGCTAc); SLC7A5 (sg3 F caccgCCCGCCCCCTCGGCCCAGCT, R aaacAGCTGGGCCGAGGGGGCGGGc; sg4 F caccgGAGGGACGGGGCCGGGCCAC, R aaacGTGGCCCGGCCCCGTCCCTCc; sg5 F caccgGTGCGTCGTCCGGCCCAGCC, R aaacGGCTGGGCCGGACGACGCACc), SLC1A2 (sg3 F caccgAGATCCTGGGCTCCTGCCAC, R aaacGTGGCAGGAGCCCAGGATCTc; sg4 F caccgGGCAGAGGAGGGACCGCGAC, R aaacGTCGCGGTCCCTCCTCTGCCc ; sg5 F caccgAAAGGAGTTGCCCGAGGCGG, R aaacCCGCCTCGGGCAACTCCTTTc), sgSLC1A3 (sg1 F caccgCACCCTCGTCTTCCCTGAAA, R aaacTTTCAGGGAAGACGAGGGTGc; sg2 F caccgAGGAAACATGCAATAATGTG, R aaacCACATTATTGCATGTTTCCTc; sg5 F caccgCTCGTAACAGTTGTACAACC, R aaacGGTTGTACAACTGTTACGAGc). sgRen targeting Renilla luciferase cDNA was used as negative-control sgRNA (GGTATAATACACCGCGCTAC)^33^ and cloned into lenti sgRNA(MS2)-zeo backbone (Addgene: 61427) or lenti sgRNA(MS2)-zeo-IRES-eGFP backbone. Codon-optimized cDNAs for human SLC1A2 and SLC1A3 were obtained from Genscript and cloned in pTO vector with or without a N-terminal Strep-HA tag.

### Transcriptomic analysis of cells under nutrient limiting conditions

HEK293T and HeLa cells were seeded in full media (DMEM Gibco, 10% FBS, antibiotics). After 24h, media was removed and, after a wash with PBS, substituted with DMEM media lacking the indicated amino acid supplemented with 10% dialyzed FBS (Gibco cat. 26400-044). Starvation media lacking a specific amino acid were prepared by complementing amino acid-free DMEM media (ie devoid of all 15 amino acids normally present, custom made by PAN biotech) with the other 14 amino acids (from individual powders, SIGMA). DMEM media reconstituted with all 15 amino acids and 10% dialyzed FBS as well as full media served as controls. For HEK293T cells the following additional conditions were tested: 100nM Torin-1 (Tocris Bioscience) in DMEM media reconstituted with all 15 amino acids and 10% dialyzed FBS; DMEM without vitamins (custom made by PAN biotech) and 10% dialyzed FBS; DMEM without glucose and pyruvate (custom made by PAN biotech) and 10% dialyzed FBS. After 16h, media was removed and cells were harvested in cold PBS. Total RNA was isolated using the Qiagen RNeasy Mini kit including a DNase I digest step. RNA-sequencing (RNA-seq) libraries were prepared using QuantSeq 3′ mRNA-Seq Library Prep Kit FWD for Illumina (Lexogen) according to the manufacturer’s protocol. Libraries were subjected to 50-bp single-end high-throughput sequencing on an Illumina HiSeq 4000 platform at the Biomedical Sequencing Facility (https://biomedical-sequencing.at/). Raw sequencing reads were demultiplexed, and after barcode, adaptor and quality trimming with cutadapt (https://cutadapt.readthedocs.io/en/stable/), quality control was performed using FastQC (http://www.bioinformatics.babraham.ac.uk/projects/fastqc/). The remaining reads were mapped to the GRCh38 (h38) human genome assembly using genomic short-read RNA-Seq aligner STAR version 2.6.1as^57^. We obtained more than 98% mapped reads in each sample with 70–80% of reads mapping to unique genomic location. Transcripts were quantified using End Sequence Analysis Toolkit (ESAT)^58^. Differential expression analysis was performed using three biological replicates with DESeq2 (1.21.21) on the basis of read counts^59^. Exploratory data analysis and visualizations were performed in R-project version 3.4.2 (Foundation for Statistical Computing, https://www.R-project.org/) with Rstudio IDE version 1.0.143, ggplot2 (3.0.0), dplyr (0.7.6), readr (1.1.1), gplots (3.0.1).

### Generation of a SLC-focused gain-of-function library

A set of single guide RNAs (sgRNAs) targeting 388 human SLC genes, with up to six sgRNA per RefSeq transcript, were generated using the Cas9 activator tool (http://sam.genome-engineering.org/database/)^34^. An additional set of 120 non-targeting sgRNAs was included by generating random 20-mers and selecting for sequences with at least three mismatches from any genomic sequence. Adapter sequences were added to the 5’ and 3’ sequences (5’prefix: TGGAAAGGACGAAACACCG, 3’suffix: GTTTTAGAGCTAGGCCAACATGAGGAT) to allow cloning by Gibson assembly in the lenti-sgRNA(MS2)_EF1A-zeo cloning backbone (addgene #61427). The oligos were synthetized as a pool by LC Sciences. Full-length oligonucleotides (66 nt) were amplified by PCR using Phusion HS Flex (NEB) and size-selected using a 2% agarose gel (Primers: SAM_ArrayF TAACTTGAAAGTATTTCGATTTCTTGGCTTTATATATCTTGTGGAAAGGAC GAAACACCG, SAM_ArrayR TTTTAACTTGCTAGGCCCTGCAGACATGGGTGATCCTCATGTTGGCCTAGCTCTAAAAC)

The vector was digested with BsmBI (NEB) for 1h at 55°C, heat inactivated for 20’ at 80°C, following by incubation with Antarctic phosphatase (NEB) for 30’ at 37°C. A 10 μl Gibson ligation reaction (NEB) was performed using 5 ng of the gel-purified inserts and 12.5 ng of the vector, incubated for 1h at 50°C and dialyzed against water for 1h at RT. The reaction was then transformed in Lucigen Endura cells and plated on two 245 mm plates. Colonies were grown at 32°C for 16-20h hours and then scraped from the plates. The plasmid was purified with the Endo-Free Mega prep kit (Qiagen) and sequenced by NGS using the approach described in Konermann *et al*^35^. The library is available on Addgene (#132561).

### Genetic screening in limiting nutrient conditions

HEK293T cells were used to generate lentiviral particles by transient transfection of lentiviral constructs (SLC-SAM library or genome-wide SAM library addgene #1000000057) and packaging plasmids psPAX2 (Addgene #12260), pMD2.G (Addgene #12259) using PolyFect (Qiagen). After 24 h the medium was changed to fresh IMDM or DMEM medium supplemented with 10% FCS and antibiotics. Viral supernatants were collected after 48 h, filtered and stored at −80 °C until further use. To generate the cells containing the SAM machinery, HEK293T cells were then infected with viral particles carrying a catalytically dead Cas9-VP64 (Addgene #61425) and MS2-P65-HSF1 constructs (Addgene #89308). For the genetic screening, dilution factors for library transduction at a MOI of 0.2-0.3 were determined by zeocin survival after transduction. For SLC SAM-based screens (Fig. 3), cells were infected with the SLC SAM library at high coverage (>1,000×) and, after selection for seven days with zeocin (250 μg ml-1), transferred in media containing modified amino acid concentrations, as listed in Suppl Fig 3a (5×10^6^ cells/condition, 2 replicates/condition). Cells in amino acid depleted conditions were passaged every 3 days, cells in full and reconstituted media when close to confluency. After 14 days, the cells were collected, washed with PBS and stored at –80C until further use. Cells grown in low methionine media for 14 days were amplified for 5 days in full media before collection. Cells grown in media lacking all amino acids for 12 days were amplified for 5 days in full media before collection. For genome-wide SAM-based screens (Fig. 5), cells were infected with the genome-wide SAM library at high coverage (1,000×) and, after selection for ten days with zeocin (250 μg ml-1), transferred in media containing either low arginine concentrations (1%) or fully reconstituted supplemented with 10% dialyzed FBS (Gibco cat. 26400-044) (50×10^6^ cells/condition, 3 replicates/condition). Cells arginine depleted media were passaged every 3 days, cells in reconstituted media when close to confluency. After 14 days, the cells were collected, washed with PBS and stored at –80C until further use. For all screens, genomic DNA was extracted using the DNAeasy kit (Qiagen) and the sgRNA cassette amplified by PCR as described in ref. 33.

### Screen analysis and hit scoring

sgRNA sequences were extracted from NGS reads, matched against the original sgRNA library index and counted using an in-house Python script. Differential abundance of individual sgRNAs was estimated using DESeq2 (1.21.21)^59^. Contrasts were performed individually for each treatment and significance was tested using either one- or two-tailed Wald tests. The sgRNAs were then sorted by p-value and aggregated into genes using GSEA (fgsea R package v1.7). To avoid false positives, only significant sgRNAs (P ≤ 0.05) were considered for enrichment, requiring also a minimum of two sgRNAs per gene. Gene enrichment significance was estimated by a permutation test using 10^8^ permutations, and p-values were corrected for multiple testing using the Benjamini–Hochberg procedure (FDR)^60^.

### Flow cytometry-based competition assay

Flow cytometry-based color competition assay were performed using either SAM-component expressing HEK293T stably transduced with the indicated sgRNA in lenti sgRNA(MS2)-zeo (Addgene: 61427, eGFP negative) and control sgRen in lenti sgRNA(MS2)-zeo-IRES-eGFP (eGFP positive) or, for cDNA-based overexpression experiments, using HEK293 FlipIN empty cells (parental) or expressing the indicated SLCs and control HEK293 FlipIN cells expressing eGFP. The cell populations were mixed in a 1:1 ratio and seeded in three replicates in full media. For cDNA-based overexpression CCA, doxycycline (1 μg/ml) was added when cells were mixed and kept for all the duration of the assay. After 24h (day 0), full media was removed and, after a wash with PBS, exchanged with the indicated media. Every 3 days, cells were detached and reseeded into fresh indicated media for the duration of the assay (12 days or as indicated) and analysed by flow cytometry.

The respective percentage of viable (FSC/SSC) eGFP-positive and eGFP-negative cells at the indicated time points was quantified by flow cytometry and ratio (eGFP-/eGFP+) were calculated. Where indicated, ratios were normalized to the ratio (eGFP-/eGFP+) at day 0.

### Immunofluorescence

SLC1A2/A3-expressing HEK293 Flip-In cells were plated on Poly-L-lysine hydrobromide (Sigma)-coated glass coverslips and induced with doxycycline (1 μg/ml) where indicated. After 24 h, cells were washed with PBS, fixed (PBS, 2% formaldehyde) and permeabilized (PBS, 0.3% saposin, 10% FBS). Slides were incubated with anti-HA (Covance, MMS-101P) and anti-AIF (Cell Signaling, 5318) antibodies (1 h, RT, PBS, 0.3% saposin, 10% FBS). After three washes slides were incubated with Alexa-Fluor-488-coupled anti-mouse (Life Technologies, A11001) and Alexa-Fluor-594-coupled anti-rabbit (Life Technologies, A11012) (1 h, RT, PBS, 0.3% saposin, 10% FBS). After DAPI staining, slides were washed three times and mounted on coverslips with ProLong Gold (Invitrogen). Images were taken with a Zeiss Laser Scanning Microscope (LSM) 700.

### Q-PCR

Total RNA was isolated using the RNeasy Mini Kit (Qiagen) including DNase I digestion step. The reverse transcription was performed using RevertAid First Strand cDNA Synthesis Kit (Thermo Scientific) using oligo dT primers. Quantitative PCR was carried out on a Rotor Gene Q (Qiagen) PCR machine using the SensiMix SYBR kit (Bioline). Results were quantified using the 2^−ΔΔCt^ method, using GAPDH as reference. The primers used were: GAPDH F: 5′-CCTGACCTGCCGTCTAGAAA-3′, R: 5′-CTCCGACGCCTGCTTCAC-3′; SLC7A3 F: 5′-AACTCGGCTTAACTCCGCCT -3′, R: 5′-CACCTCGCCAGCTAGGACAT -3′.

### Immunoblot

Cells were lysed in E1A lysis buffer (50 mM HEPES, 250 mM NaCl, 5 mM EDTA, 1% NP-40, pH 7.4) supplemented with EDTA-free protease inhibitor cocktail (Roche, 1 tablet per 50 ml) and Halt phosphatase inhibitor cocktail (Thermo Fisher Scientific) for 10 min on ice. Lysates were cleared by centrifugation at 13,000 rpm, 10 min, 4 °C and normalized by BCA (Thermo Fisher Scientific) using BSA as standard. Where indicated, cleared lysate was incubated without or with PNGase F (NEB, 1 μl per 40 μl lysate) for 30 min at 37 °C. Typically, 10-20 μg protein per sample was resolved by SDS–PAGE and blotted to nitrocellulose membranes. Membranes were blocked with 5% non-fat dry milk in TBST, probed with the indicated antibodies and detected with horseradish-peroxidase-conjugated secondary antibodies.

### Metabolomics

The HEK293 Jump-In TREx SLC7A3 cell line was plated at sub-confluent density in cell-culture treated 6-well plate in normal growth medium with or without 1μg/ml doxycycline (six replicates/condition, 750000 cells/well). Cells were incubated overnight at 37°C, 21% O2 in a humidified incubator. The following day, cells were washed gently two times with 500 μl PBS (room temperature). The plate was transferred to ice and 1500 μl/well of 80:20 ice-cold MeOH:H_2_O solution was added, cells were scraped and transferred to pre-cooled Eppendorf tubes. Samples were shortly vortexed, snap frozen in liquid nitrogen, thawed on ice and centrifuged at 20800g at 4°C for 10 min. Supernatant was transferred into a HPLC vial and evaporated using a nitrogen evaporator. Evaporated samples were reconstituted in 50 μL of 80:20 MeOH:H2O and stored at -80°C until analysis. The targeted metabolomics was performed using the Biocrates Absolute IDQ p180 kit. The kit provides (semi)-quantitative analysis of 184 metabolites. Sample preparation was performed according to the User Manual provided by the manufacturer. Briefly, 10 μL of internal standard was loaded onto the 96-well kit plate, dried under nitrogen and followed by loading 10 μL of samples, blanks, calibration standards, and quality control samples. After an additional drying step, derivatization reagent was added, and samples were incubated for 20 min at room temperature. The 96-well plate was dried, and the extraction solvent was added. Supernatants were collected in a new 96-well kit plate by centrifugation, diluted and then analysed using a Water Acquity UHPLC system and Waters Xevo TQ-MS mass spectrometer. Each sample was analyzed in using two methods: 1) Metabolites were measured with LC-MS/MS method (positive ionization mode, sample injected using a HPLC column), 2) Lipids were measured with FIA-MS/MS method (positive mode, sample injected without a HPLC column). The UHPLC method files KIT2_LC_8015.IPR (.w2200, .wvhp) and KIT3-FIA_8015.IPR (.w2200, .wvhp) are part of the kit provided by Biocrates. The acquired LC-MS/MS data was processed with Waters MassLynx V4.1 software TargetLynx module using the processing method KIT2_WatersQuan_8015.mdb provided with the kit by Biocrates. The obtained results were imported in Biocrates METIDQ software where further validation of calibration standards and quality control sample accuracy as well as internal standard intensity were carried out. The acquired raw FIA-MS/MS data were directly imported in Biocrates METIDQ software where it was processed and validated. The final results were exported as tsv files. Conditions were compared using Welch’s t-test, p-value was subsequently corrected for multiple testing according to the Benjamini and Hochberg procedure^60^.

## References

1. Torrence, M. E. & Manning, B. D. Nutrient Sensing in Cancer. Annu. Rev. Cancer Biol. 2, 251–269 (2018).

2. Palm, W. & Thompson, C. B. Nutrient acquisition strategies of mammalian cells. Nature 546, 234–242 (2017).

3. Sullivan, M. R. & Vander Heiden, M. G. Determinants of nutrient limitation in cancer. Crit. Rev. Biochem. Mol. Biol. 1–15 (2019) doi:10.1080/10409238.2019.1611733.

4. Finicle, B. T., Jayashankar, V. & Edinger, A. L. Nutrient scavenging in cancer. Nat. Rev. Cancer 18, 619–633 (2018).

5. Gardner, D. K. Changes in requirements and utilization of nutrients during mammalian preimplantation embryo development and their significance in embryo culture. Theriogenology 49, 83–102 (1998).

6. Kilberg, M. S., Terada, N. & Shan, J. Influence of Amino Acid Metabolism on Embryonic Stem Cell Function and Differentiation. Adv. Nutr. 7, 780S–789S (2016).

7. Lu, V., Roy, I. J. & Teitell, M. A. Nutrients in the fate of pluripotent stem cells. Cell Metab. 33, 2108–2121 (2021).

8. Kedia-Mehta, N. & Finlay, D. K. Competition for nutrients and its role in controlling immune responses. Nat. Commun. 10, 2123 (2019).

9. Altea-Manzano, P., Cuadros, A. M., Broadfield, L. A. & Fendt, S.-M. Nutrient metabolism and cancer in the in vivo context: a metabolic game of give and take. EMBO Rep. 21, e50635 (2020).

10. Hosios, A. M. et al. Amino Acids Rather than Glucose Account for the Majority of Cell Mass in Proliferating Mammalian Cells. Dev. Cell 36, 540–549 (2016).

11. Bento, C. F. et al. Mammalian Autophagy: How Does It Work? Annu. Rev. Biochem. 85, 685–713 (2016).

12. Fung, M. K. L. & Chan, G. C.-F. Drug-induced amino acid deprivation as strategy for cancer therapy. J. Hematol. Oncol.J Hematol Oncol 10, 144 (2017).

13. Cramer, S. L. et al. Systemic depletion of L-cyst(e)ine with cyst(e)inase increases reactive oxygen species and suppresses tumor growth. Nat. Med. 23, 120–127 (2017).

14. Broome, J. D. Evidence that the L -Asparaginase Activity of Guinea Pig Serum is responsible for its Antilymphoma Effects. Nature 191, 1114 (1961).

15. Butler, M., Meer, L. T. van der & Leeuwen, F. N. van. Amino Acid Depletion Therapies: Starving Cancer Cells to Death. Trends Endocrinol. Metab. 32, 367–381 (2021).

16. Feun, L., Kuo, M. & Savaraj, N. Arginine deprivation in cancer therapy. Curr. Opin. Clin. Nutr. Metab. Care 18, 78–82 (2015).

17. Nicklin, P. et al. Bidirectional Transport of Amino Acids Regulates mTOR and Autophagy. Cell 136, 521–534 (2009).

18. van Geldermalsen, M. et al. ASCT2/SLC1A5 controls glutamine uptake and tumour growth in triple-negative basal-like breast cancer. Oncogene 35, 3201–3208 (2016).

19. Cormerais, Y. et al. The glutamine transporter ASCT2 (SLC1A5) promotes tumor growth independently of the amino acid transporter LAT1 (SLC7A5). J. Biol. Chem. 293, 2877–2887 (2018).

20. Karunakaran, S. et al. SLC6A14 (ATB0,+) protein, a highly concentrative and broad specific amino acid transporter, is a novel and effective drug target for treatment of estrogen receptor-positive breast cancer. J. Biol. Chem. 286, 31830–31838 (2011).

21. Babu, E. et al. Deletion of the amino acid transporter Slc6a14 suppresses tumour growth in spontaneous mouse models of breast cancer. Biochem. J. 469, 17–23 (2015).

22. Cormerais, Y. et al. Genetic Disruption of the Multifunctional CD98/LAT1 Complex Demonstrates the Key Role of Essential Amino Acid Transport in the Control of mTORC1 and Tumor Growth. Cancer Res. 76, 4481–4492 (2016).

23. Wellberg, E. A. et al. The glucose transporter GLUT1 is required for ErbB2-induced mammary tumorigenesis. Breast Cancer Res. BCR 18, 131 (2016).

24. Hediger, M. A., Clémençon, B., Burrier, R. E. & Bruford, E. A. The ABCs of membrane transporters in health and disease (SLC series): Introduction. Mol. Aspects Med. 34, 95–107 (2013).

25. César-Razquin, A. et al. A Call for Systematic Research on Solute Carriers. Cell 162, 478–487 (2015).

26. O’Hagan, S., Wright Muelas, M., Day, P. J., Lundberg, E. & Kell, D. B. GeneGini: Assessment via the Gini Coefficient of Reference “Housekeeping” Genes and Diverse Human Transporter Expression Profiles. Cell Syst. 6, 230–244 (2018).

27. César-Razquin, A. et al. In silico Prioritization of Transporter–Drug Relationships From Drug Sensitivity Screens. Front. Pharmacol. 9, 1011 (2018).

28. Kilberg, M. S., Shan, J. & Su, N. ATF4-dependent transcription mediates signaling of amino acid limitation. Trends Endocrinol. Metab. TEM 20, 436–443 (2009).

29. Bacci, M. et al. Reprogramming of Amino Acid Transporters to Support Aspartate and Glutamate Dependency Sustains Endocrine Resistance in Breast Cancer. Cell Rep. 28, 104–118.e8 (2019).

30. Carette, J. E. et al. Haploid genetic screens in human cells identify host factors used by pathogens. Science 326, 1231–1235 (2009).

31. Hsu, P. D., Lander, E. S. & Zhang, F. Development and Applications of CRISPR-Cas9 for Genome Engineering. Cell 157, 1262–1278 (2014).

32. Reczek, C. R. et al. A CRISPR screen identifies a pathway required for paraquat-induced cell death. Nat. Chem. Biol. 13, 1274–1279 (2017).

33. Fauster, A. et al. Systematic genetic mapping of necroptosis identifies SLC39A7 as modulator of death receptor trafficking. Cell Death Differ. 26, 1138–1155 (2018).

34. Chavez, A. et al. Comparison of Cas9 activators in multiple species. Nat. Methods 13, 563–567 (2016).

35. Konermann, S. et al. Genome-scale transcriptional activation by an engineered CRISPR-Cas9 complex. Nature 517, 583–8 (2014).

36. Gilbert, L. A. et al. Genome-Scale CRISPR-Mediated Control of Gene Repression and Activation. Cell (2014) doi:10.1016/j.cell.2014.09.029.

37. Meixner, E. et al. A substrate-based ontology for human solute carriers. Mol. Syst. Biol. 16, (2020).

38. Thoreen, C. C. et al. An ATP-competitive Mammalian Target of Rapamycin Inhibitor Reveals Rapamycin-resistant Functions of mTORC1. J. Biol. Chem. 284, 8023–8032 (2009).

39. Labuschagne, C. F., van den Broek, N. J. F., Mackay, G. M., Vousden, K. H. & Maddocks, O. D. K. Serine, but not glycine, supports one-carbon metabolism and proliferation of cancer cells. Cell Rep. 7, 1248–1258 (2014).

40. Locasale, J. W. Serine, glycine and one-carbon units: cancer metabolism in full circle. Nat. Rev. Cancer 13, 572–583 (2013).

41. Mazor, K. M. et al. Effects of single amino acid deficiency on mRNA translation are markedly different for methionine versus leucine. Sci. Rep. 8, (2018).

42. Han, H. et al. TRRUST v2: An expanded reference database of human and mouse transcriptional regulatory interactions. Nucleic Acids Res. 46, D380–D386 (2018).

43. Kandasamy, P., Gyimesi, G., Kanai, Y. & Hediger, M. A. Amino acid transporters revisited: New views in health and disease. Trends Biochem. Sci. (2018) doi:10.1016/j.tibs.2018.05.003.

44. Park, Y., Reyna-Neyra, A., Philippe, L. & Thoreen, C. C. mTORC1 Balances Cellular Amino Acid Supply with Demand for Protein Synthesis through Post-transcriptional Control of ATF4. Cell Rep. 19, 1083–1090 (2017).

45. Fotiadis, D., Kanai, Y. & Palacín, M. The SLC3 and SLC7 families of amino acid transporters. Mol. Aspects Med. 34, 139–158 (2013).

46. Tajan, M. et al. A Role for p53 in the Adaptation to Glutamine Starvation through the Expression of SLC1A3. Cell Metab. 28, 721–736.e6 (2018).

47. Alkan, H. F. et al. Cytosolic Aspartate Availability Determines Cell Survival When Glutamine Is Limiting. Cell Metab. 28, 706–720.e6 (2018).

48. Palmieri, F. The mitochondrial transporter family SLC25: Identification, properties and physiopathology. Mol. Aspects Med. 34, 465–484 (2013).

49. Vander Heiden, M. G. & DeBerardinis, R. J. Understanding the Intersections between Metabolism and Cancer Biology. Cell 168, 657–669 (2017).

50. Chapman, N. M., Boothby, M. R. & Chi, H. Metabolic coordination of T cell quiescence and activation. Nat. Rev. Immunol. 20, 55–70 (2020).

51. Lyssiotis, C. A. & Kimmelman, A. C. Metabolic Interactions in the Tumor Microenvironment. Trends Cell Biol. 27, 863–875 (2017).

52. Birsoy, K. et al. Metabolic determinants of cancer cell sensitivity to glucose limitation and biguanides. Nature 508, 108–112 (2014).

53. Gonzalez-Menendez, P. et al. GLUT1 protects prostate cancer cells from glucose deprivation-induced oxidative stress. Redox Biol. 17, 112–127 (2018).

54. Kilberg, M. S., Pan, Y.-X., Chen, H. & Leung-Pineda, V. Nutritional Control of Gene Expression: How Mammalian Cells Respond to Amino Acid Limitation. Annu. Rev. Nutr. 25, 59–85 (2005).

55. McCracken, A. N. & Edinger, A. L. Targeting cancer metabolism at the plasma membrane by limiting amino acid access through SLC6A14. Biochem. J. 470, e17–e19 (2015).

56. Superti-Furga, G. et al. The RESOLUTE consortium: unlocking SLC transporters for drug discovery. Nat. Rev. Drug Discov. (2020) doi:10.1038/d41573-020-00056-6.

57. Dobin, A. et al. STAR: ultrafast universal RNA-seq aligner. Bioinformatics 29, 15–21 (2013).

58. Derr, A. et al. End Sequence Analysis Toolkit (ESAT) expands the extractable information from single-cell RNA-seq data. Genome Res. 26, 1397–1410 (2016).

59. Love, M. I., Huber, W. & Anders, S. Moderated estimation of fold change and dispersion for RNA-seq data with DESeq2. Genome Biol. 15, 550 (2014).

60. Benjamini, Y. & Hochberg, Y. Controlling the False Discovery Rate: A Practical and Powerful Approach to Multiple Testing. J. R. Stat. Soc. Ser. B Methodol. 57, 289–300 (1995).

