## Supplementary Figures for "Gain-of-function genetic screens in human cells identify SLC transporters overcoming environmental nutrient restrictions"

**Figure S1: a.** Read counts for the transcriptomics samples in HEK293T and HeLa cells. Box plots showing the median and 25/75 percentile are shown. Samples excluded from further analysis due to low counts are shown in dark grey. **b.** PCA plot comparing all samples used in further analysis from both HEK293T and HeLa cells. **c.** Enrichment analysis for target genes of transcription factors in the HeLa dataset, as determined by TRRUST v2. **d.** Heatmap showing differential expression of SLC transporters in the listed conditions compared to reconstituted media in HeLa cells.

**Figure S2: a.** Schematic representation of the SAM technology for transcriptional activation. **b.** Read counts for SLC SAM library sgRNAs in purified plasmid sample. **c.** Representative images of HEK293T cells treated for 7 days with either full media and dialysed FBS (top) or media with 5% of the original arginine concentration and dialysed FBS. The bar corresponds to 500 µM. **d.** Expression of SLC7A3 in the indicated sgRNA-expressing SAM-HEK293T cells assessed by Q-PCR.

**Figure S3: a.** Screening conditions applied to SLC-SAM-infected HEK293T cells. **b.** Plots showing the normalized counts of sgRNAs targeting the listed SLC-targeting across the conditions tested. **c.** Overview of the significantly enriched SLCs for each of the tested conditions at a FDR < 0.05. The circle size reflects the absolute log2 fold change (aLFC) above the fully reconstituted media, while the distance from the center reflects the statistical significance of the effect observed.

**Figure S4: a-b.** Western blot showing expression of endogenous SLC1A2 (a) and SLC1A3 SAM-HEK293T cells expressing the indicated sgRNA. Where indicated, cell lysates were treated with PNGase. **c-e.**Western blot showing expression of Strep-HA-tagged (c) or untagged SLC1A2, SLC1A3 and GFP constructs in HEK293 Flip In cells with (+) or without(-) doxycycline-induction. Where indicated, cell lysates were treated with PNGase. **f-g**. Confocal microscopy images of

HEK293 Flip In cells expressing Strep-HA-tagged SLC1A2 (**f**) or SLC1A3 (**g**) with (+) or without (-) doxycycline-induction. AIF staining marks mitochondria.

**Figure S5: a.** Targeted metabolomics analysis of doxycycline-induced HEK293 Jump In cells overexpressing SLC7A3, compared to non-induced cells, show enrichment for transporter substrates and related compounds. Only results for amino acids and biogenic amines are shown. Mean and SEM of one experiment, performed in six technical replicates, are shown. AA. Amino acids, AC, acylcarnitines, BA, biogenic amines. **b.** Concentration of the significantly enriched or depleted metabolites in HEK293 Jump In cells overexpressing SLC7A3 induced (+) or not (-) with doxycyline. **c.** Western blot image showing doxycline-dependent expression of HA-tagged SLC7A3 in Jump In TREx cells. Cyclophilin B is shown as loading control.

### Supplementary Figure 1

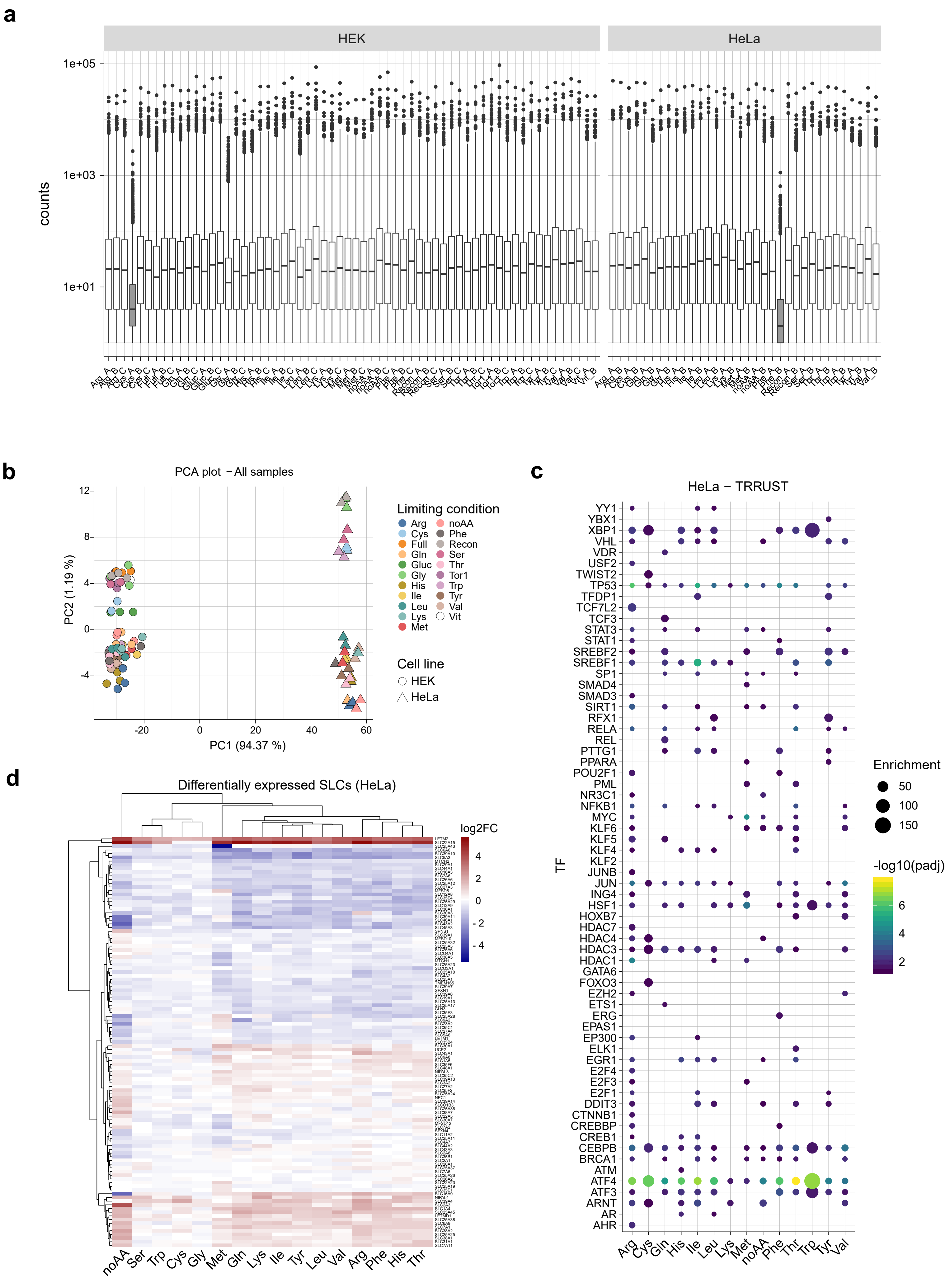

#### Supplementary Figure 2

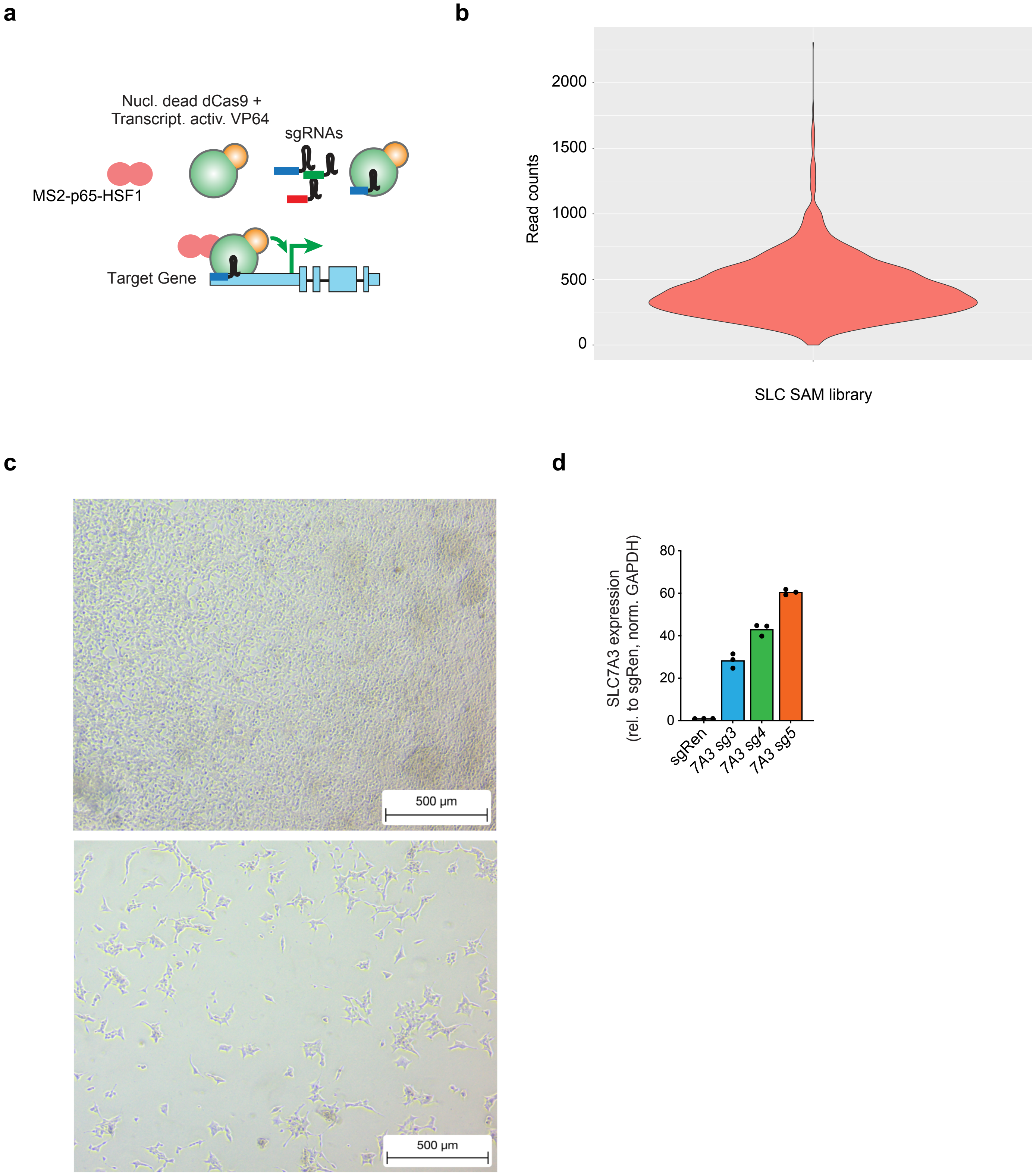

#### Supplementary Figure 3

**a**

| AA | (% of DMEM) |
|---|---|
| Glutamine | 0.1 |
| Arginine | 1 |
| Cystine | 2 |
| Histidine | 1 |
| Isoleucine | 1 |
| Leucine | 1 |
| Lysine | 1 |
| Methionine | 2 |
| Phenylalanine | 1 |
| Threonine | 1 |
| Tryptophan | 1 |
| Tyrosine | 1 |
| Valine | 1 |
| Recon + Dialyzed FBS | |
| No AAs + Dialyzed FBS | |
| Full DMEM + FBS | |
| Day 0 (T0) | |

**b**

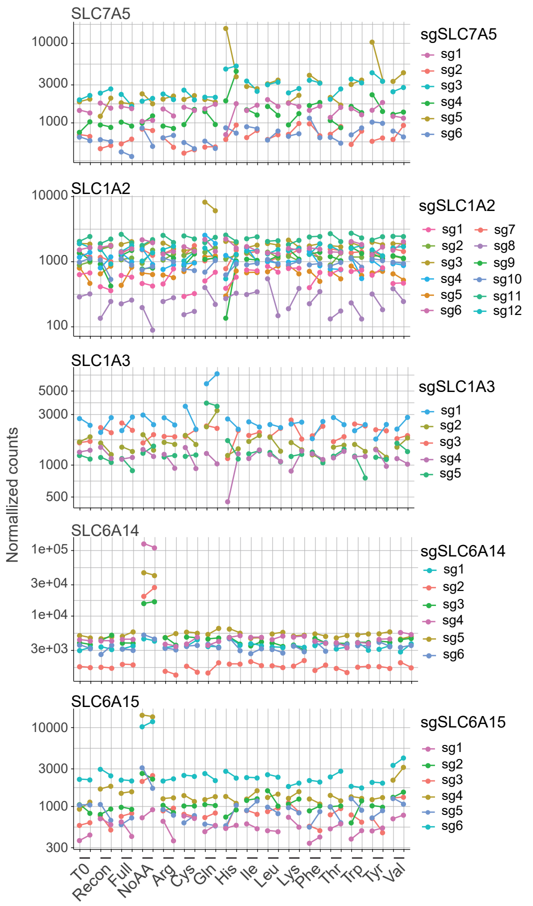


**c**

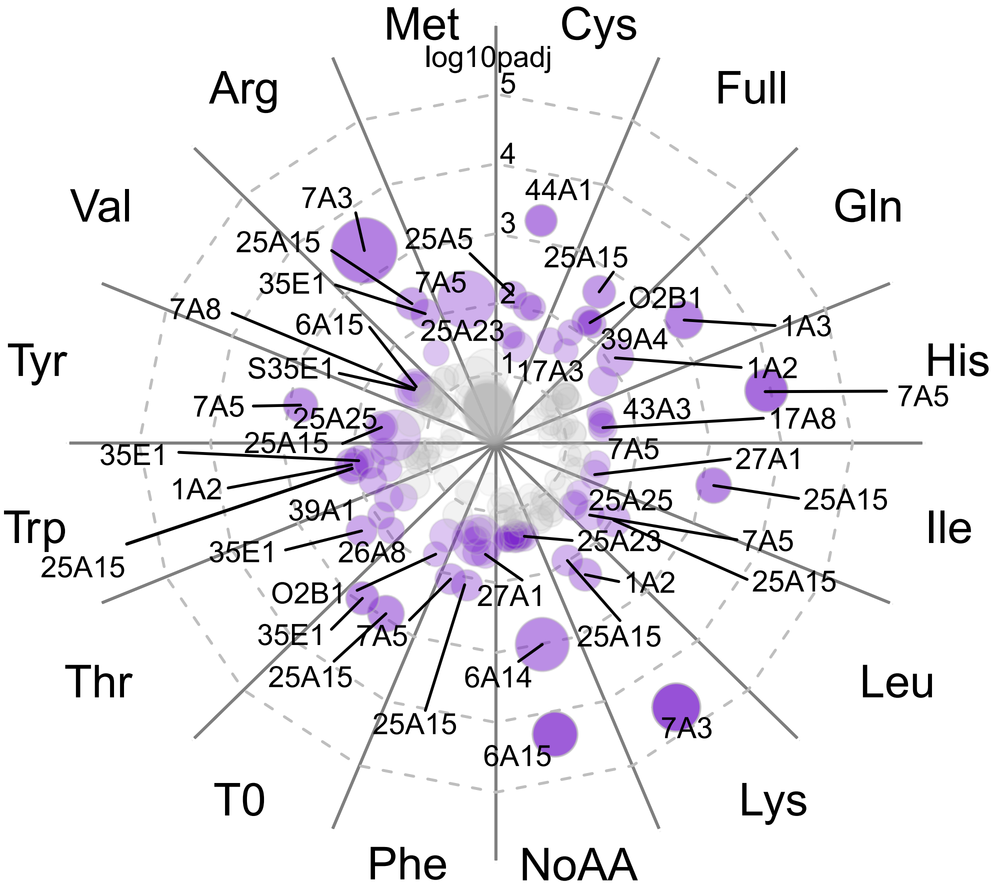

**Supplementary Figure 4**

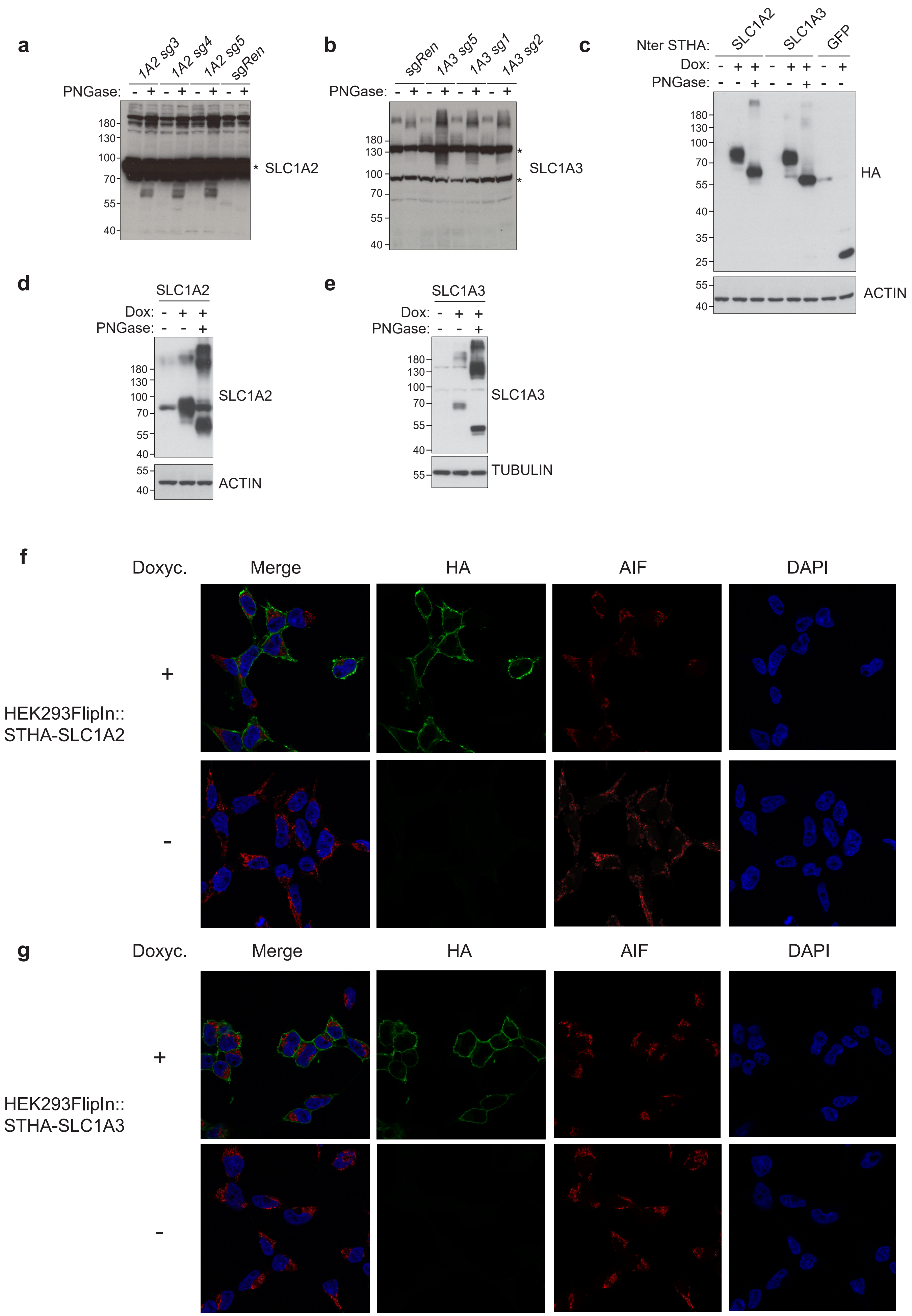

#### Supplementary Figure 5

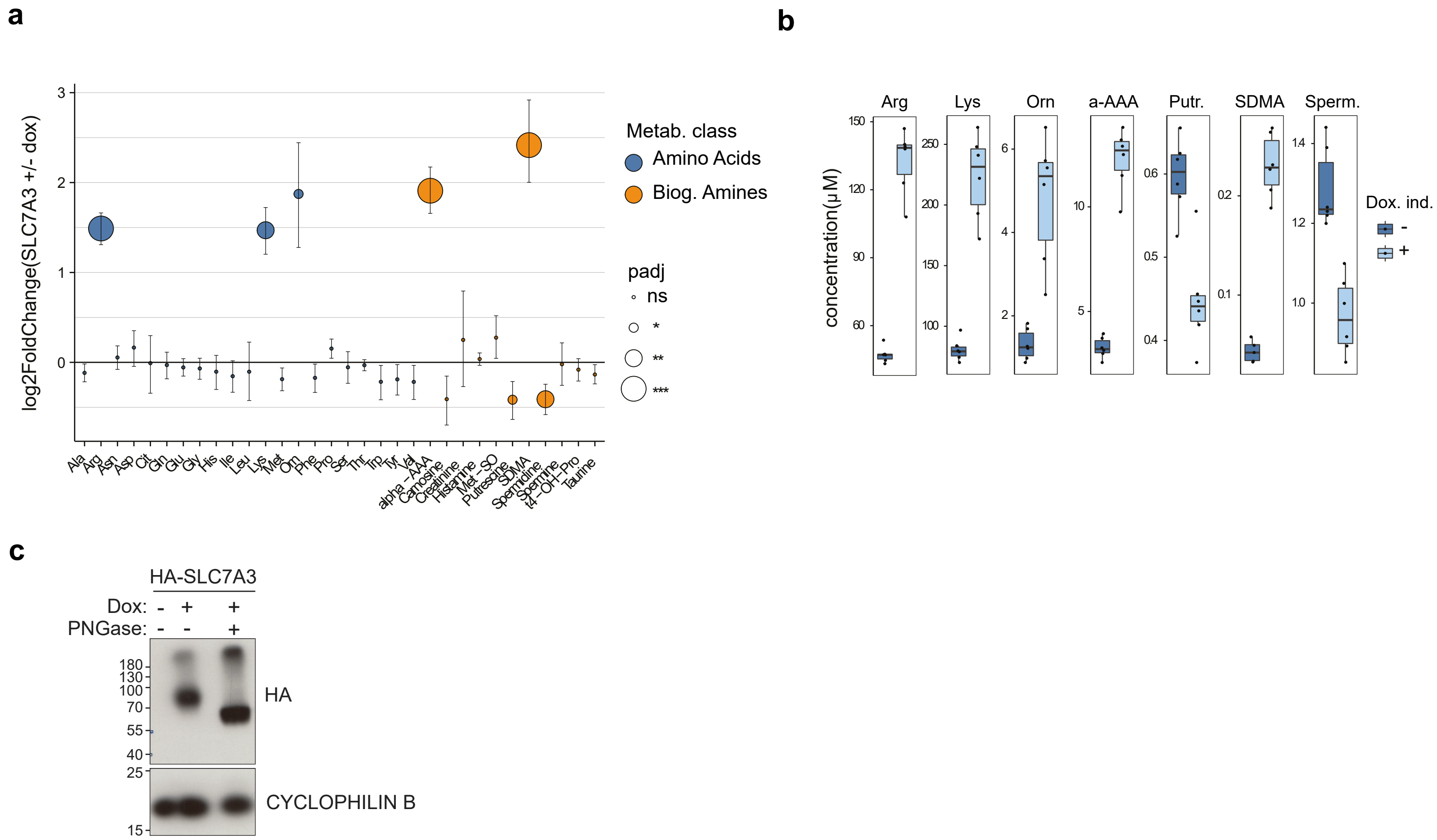
